# Elucidating an anterior cingulate circuit for self-initiated actions and rescue of Parkinsonian akinesia

**DOI:** 10.64898/2025.12.01.691607

**Authors:** Francesco P. Ulloa Severino, Bryan Lu, Jiwon Kim, Alexander Friedman, Marina Roshchina, Sarah A. Johnson, Konstantin I. Bakhurin, Cagla Eroglu, Henry H. Yin

**Affiliations:** Cajal Institute, CSIC, Madrid 28002. Spain; Cajal Neuroscience Center, CSIC, Campus de Alcalá de Henares, Madrid 28805. Spain; Department of Psychology and Neuroscience, Duke University; Durham, USA; Department of Cell Biology, Duke University Medical Center, Durham, USA; Department of Neurobiology, Duke University Medical Center, Durham, USA; Duke Institute for Brain Sciences (DIBS), Durham, USA; Howard Hughes Medical Institute, Duke University, Durham, USA; Aligning Science Across Parkinson’s (ASAP) Collaborative Research Network, Chevy 16 Chase, MD

## Abstract

Dopamine (DA) depletion is known to result in Parkinsonian symptoms such as the inability to initiate movements (akinesia). While Parkinsonian akinesia is traditionally associated with reduced DA signaling in the striatum, the contribution of cortical regions that also receive DA projections and project to the striatum remains unclear. Here, we identify a previously unexplored cortical circuit involving D1-like dopamine receptor-expressing neurons in the anterior cingulate cortex (ACC) that is critical for initiating goal-directed movements. We find that a selective activation of ACC-D1+ neurons can flexibly drive targeted movement and locomotion even in akinetic mice after dopamine depletion or receptor antagonism. These findings uncover a cortical mechanism for movement initiation and offer promising new therapeutic targets for treating Parkinsonian akinesia.

## Introduction

Akinesia, the inability to initiate voluntary movement, is one of the most debilitating symptoms of Parkinson’s disease. Parkinson’s disease has traditionally been attributed to dopaminergic degeneration in the midbrain, which disrupts basal ganglia function and leads to motor impairments ^1,2^. The cerebral cortex, which provides the major inputs to the basal ganglia, is a key component of cortico-basal ganglia-thalamic loops ^3,4^. However, while the role of the basal ganglia in Parkinson’s disease is well-established, the role of cortical regions in motor initiation and Parkinsonian deficits remains underexplored.

The anterior cingulate cortex (ACC), a major prefrontal region, is implicated in diverse functions, including effort regulation, approach behavior, and cognitive conflict ^5–10^. Within the cortico-basal ganglia framework, the ACC forms part of the associative loop, sending robust projections to the dorsomedial striatum (DMS), a region critical for goal-directed actions and locomotion ^3,11–13^. Stimulation of the ACC can evoke movements, though typically with higher thresholds than classic motor cortices ^14^. Lesions to this region can result in reduced and slowed movements, hypometria and hypokinesia ^15,16^. Notably, the ACC receives significant dopaminergic innervation, and DA signaling within this region modulates effort-based decision-making and approach behavior ^17^. However, previous studies have not investigated the contributions of specific neuronal populations within the ACC, particularly those expressing dopamine receptors (D1+ and D2+).

In this study, using a combination of optogenetics, in vivo calcium imaging, and DA sensors in freely moving mice, we examined the functional roles of ACC neuronal populations in action initiation. Our experiments demonstrate that selective activation of ACC-D1+ neurons induces approach behavior in normal mice and restores movement in mice with complete akinesia, induced either pharmacologically with DA antagonists or through DA depletion. These findings reveal a previously unrecognized cortical circuit mechanism for volitional action and open new avenues for targeted, circuit-based interventions in Parkinson’s disease.

## Results

### Closed-loop optogenetic stimulation system and video tracking to study of whole-body targeted approach behaviors

The ACC receives dopaminergic projections mostly from the ventral tegmental area, whose activity is involved in controlling target approach or retreat ^18^. Both D1 and D2 receptors are found in the ACC, though in distinct neuronal populations ^19^. We selectively targeted ACC-D1+ and ACC-D2+ neurons that project to the DMS for closed-loop optogenetic stimulation. Transgenic mice expressing Cre in D1R+ neurons (Drd1-Cre) or D2R+ neurons (Drd2-Cre) were injected with a retrograde AAV expressing a Cre-dependent ChR2 into the DMS (“ACC-D1+” and “ACC-D2+” groups) (Fig. 1a,b). An additional control group was prepared by injecting wild-type mice (C57Bl-6J) with a retrograde AAV expressing Cre into the DMS and an AAV expressing a Cre-dependent channelrhodopsin-2 (ChR2) into the ACC to target all the ACC neurons projecting to the DMS (“ACC” group) (Supplementary Fig.1a). All groups had optic fibers bilaterally implanted in the ACC.

**Fig. 1.**
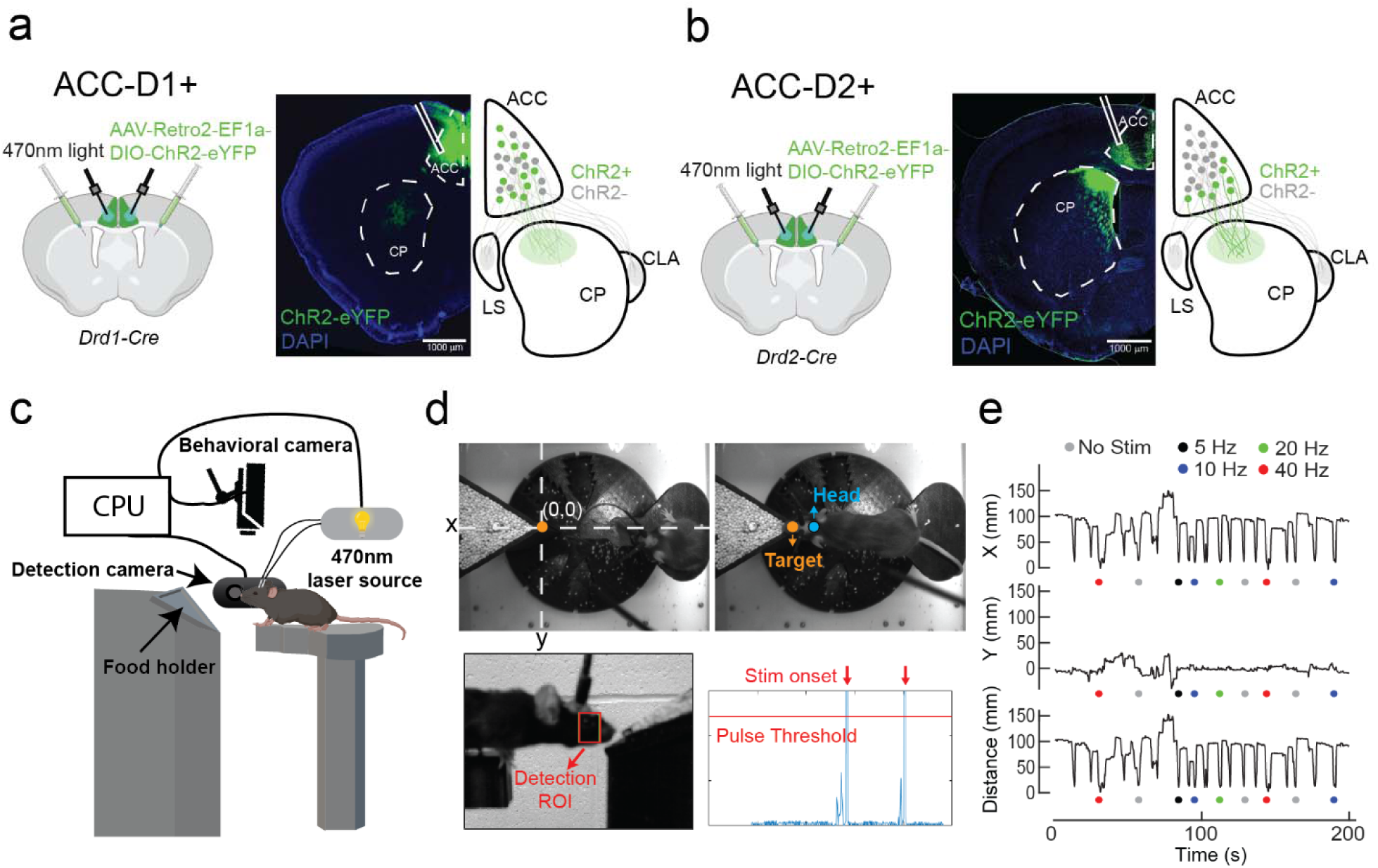
Closed-loop system for cell-type-specific and circuit-specific study of whole-body targeted movements. (**a**) Cre-dependent ChR2 expression in D1R+ neurons in the ACC (ACC-D1+). Retrograde labeling of ACC neurons that project to the DMS. (**b**) Cre-dependent ChR2 expression in D2R+ neurons in the ACC (ACC-D2+). Retrograde labeling of ACC neurons that project to the DMS. LS, lateral septum, CLA, claustrum. (**c**) Illustration of the behavioral setup used for the target approach task. The mouse stands on a platform, and food pellets or seeds are placed on a separate platform at a fixed distance (∼25 mm). One camera on one side tracks the mouse’s approach to the platform, which is used for closed-loop optogenetic stimulation. Another camera from the side records the movement. (**d**) Illustration of behavioral task and movement quantification, and an example of closed-loop optogenetic stimulation with ROI detection and triggering light stimulation upon target approach. (**e**) Euclidean variables of the head referenced to the target and extracted distance from the target, which shows the major contribution of X over Y. Colored dots represent the randomly selected stimulation parameters.

We designed a reward approach task that allows precise quantification of the behavior and closed-loop optogenetic stimulation to examine the contribution of the ACC. The mouse stands on a platform, which is separated from another platform holding food rewards (target) by a small gap (Fig. 1c and Supplementary Fig.1b). A “behavior” camera recorded the mouse movement (Fig. 1d, top) and a detection camera, with a side view, was programmed to trigger closed-loop laser stimulation during approach behavior (Fig. 1d, bottom). Mice were trained for 2-3 days until they had consistent target approaches for 10-15 minutes before testing using optogenetic stimulation. Random sampling without replacement method was used to select among the five stimulation parameters, either frequency or duration, in different sessions. Video data were analyzed using DeepLabCut ^20^ to extract head X and Y coordinates referenced to the target and calculate the distance from the target, showing no lateral movements as indicated by the small contribution of movement in Y (Fig. 1e and Supplementary Fig.1c). No differences among groups were observed in the reach frequency along the entire session (Supplementary Fig.1d).

### ACC-D1+ neurons selectively promote goal-directed approach behavior

Each approach event is characterized by an initial increase in forward velocity that marks the beginning of the approach to the target. During the approach, the distance from the mouse head to the target decreases until the target is reached (Fig. 2a, b).

**Fig. 2.**
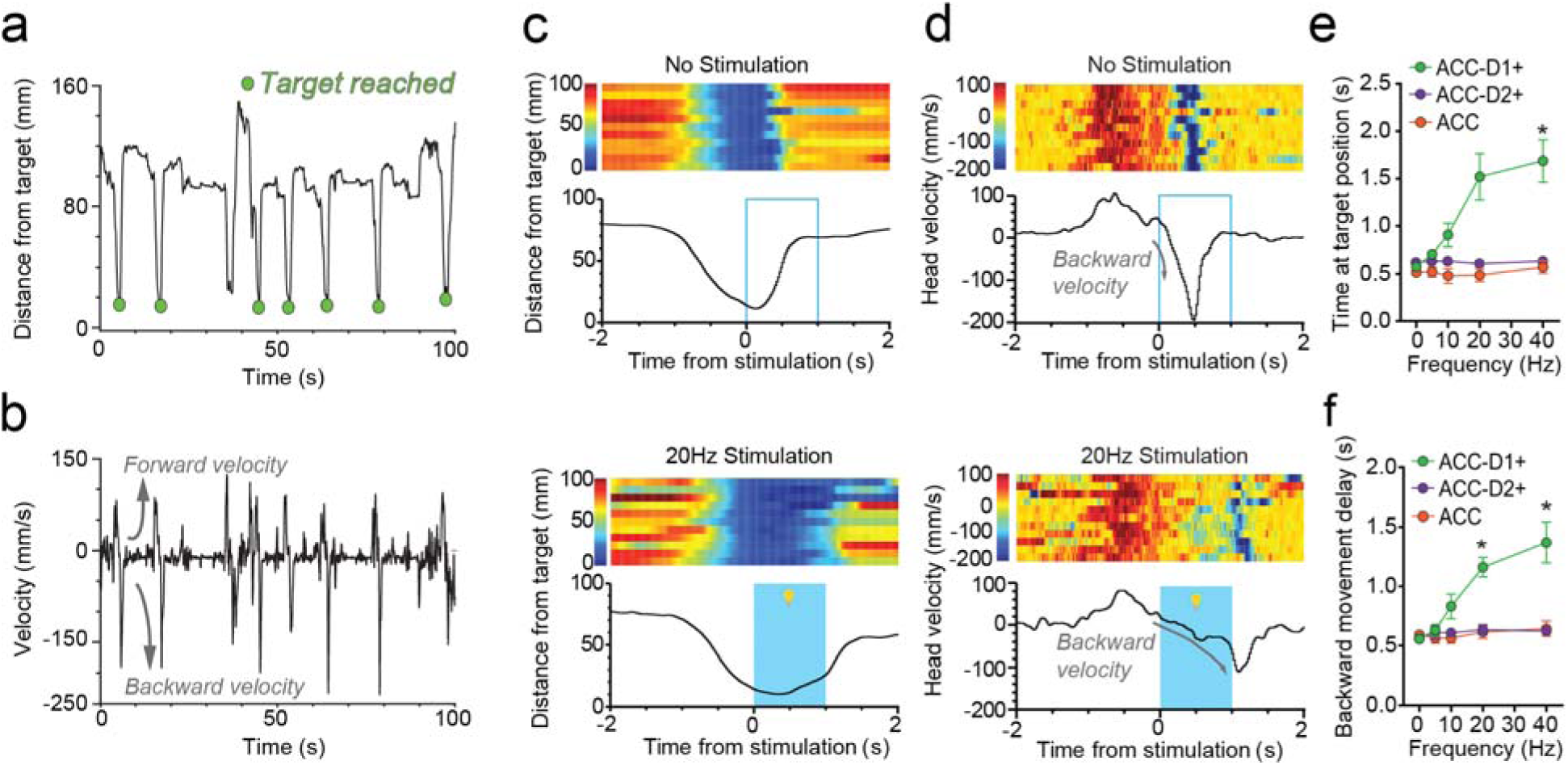
Activation of ACC-D1+ neurons generates fixation at the target. (**a**) Representative example of distance from target trace and detection of when the target has been reached by the mouse (green). (**b**) Example of a velocity trace with positive values representing forward velocity and negative values representing backward velocity. (**c**) (Top) Representative distance to target measure from a Drd1-Cre mouse during reaching in the absence of photo-stimulation. (Bottom) Distance measure from the same mouse on stimulation trials (1 s, 20 Hz). Stimulation was triggered by the target approach. (**d**) Top, representative distance to target measure from a Drd1-Cre mouse during reaching in the absence of photo-stimulation. Bottom, distance measure from the same mouse on stimulation trials (1 s, 20 Hz). (**e**) Stimulation increased time spent at the target in a frequency-dependent manner. ACC neurons (n = 7 mice) and ACC-D2+ projection (n = 4 mice) stimulations do not affect the mouse movement, whereas stimulations of ACC-D1+ projection (n = 10 mice) show a frequency-dependent effect on time spent at the target position. 2-way RM ANOVA: significant effect of frequency, F (2.114, 38.05) = 4.616, p = 0.0146; significant effect of projection-specificity, F (2, 18) = 8.093, p = 0.0031; Significant interaction, F (8, 72) = 5.432, p < 0.0001. Post hoc tests show a significant difference between ACC and ACC-D1+ groups at 40 Hz (p = 0.0326). (**f**) Stimulation caused a delay in returning to the starting position. ACC projection (n = 7 mice) and ACC-D2+ projection (n = 4 mice) stimulations do not affect the mouse backward movement, whereas stimulations of ACC-D1+ projection (n = 10 mice) show a frequency-dependent effect on peak backward velocity time. 2-way RM ANOVA: significant effect of frequency F (2.078, 37.41) = 5.852, p = 0.0056; significant effect of neuronal population, F (2, 18) = 9.082, p = 0.0019; significant interaction, F (8, 72) = 4.738, p = 0.0001. Post hoc tests show a significant difference between ACC and ACC-D1+ groups at 20 Hz (p = 0.02) and ACC-D2+ and ACC-D1+ groups at 40 Hz (p = 0.04). For all the data the degrees of freedom were adjusted using the Geisser-Greenhouse correction.

Once the mouse collects the pellet reward, it retreats from the target to the starting position. This backward retreating behavior is characterized by a large increase in backward velocity (Fig. 1c, d, top). To determine the causal role of the ACC, we performed closed-loop optogenetic stimulation. First, we stimulated ACC-D1+ neurons as soon as the mouse reached the target. Such stimulation prolonged the time spent at the target and delayed the retreat movement until the end of the stimulation (Fig. 1c, d, bottom). Such “fixation” at the target position and delay in returning to the start position are frequency-dependent (Fig. 1e,f). It can only be observed with optogenetic stimulation of ACC-D1+ projections (Movie 1). Activation of ACC-D2+ neurons or non-selective activation of all ACC neurons that project to the DMS did not produce clear effects (Fig. 1e, f, and Supplementary Fig.1e-g). Moreover, unilateral stimulation of ACC-D1+ projections is sufficient to cause a significant frequency-dependent increase in the time spent at the target position and backward movement delay (Supplementary Fig.2a-c).

Stimulation of ACC-D1+ projections also produced target “overshooting” when the stimulation frequency was high (40 Hz; Supplementary Fig.3a, b). Mice reached beyond the target location as a result of stimulation, but such overshooting was accompanied by a reduction in backward velocity during subsequent retreat behavior (Supplementary Fig.3c, d). These findings suggest that activation of ACC-D1+ projections not only results in fixation at the target but can even push the mouse further forward beyond the target when the stimulation is strong enough.

### Activation of ACC-D1+ neurons generates flexible targeted movements depending on the behavioral state

To test if the induction of overshooting depends on the stimulation parameters like frequency or pulse number, we fixed the frequency of stimulation at 20 Hz and varied stimulation duration/pulse number (0.5-3s). We found a strong linear relationship between time spent at the target position and stimulation duration. Longer stimulation durations also resulted in target overshooting (Fig. 3a, left; Fig. 3b, c). On the other hand, backward velocity in the retreat behavior was not affected, but retreat behavior (peak of backward velocity) was delayed in proportion to stimulation duration (Fig. 3d, e).

**Fig. 3.**
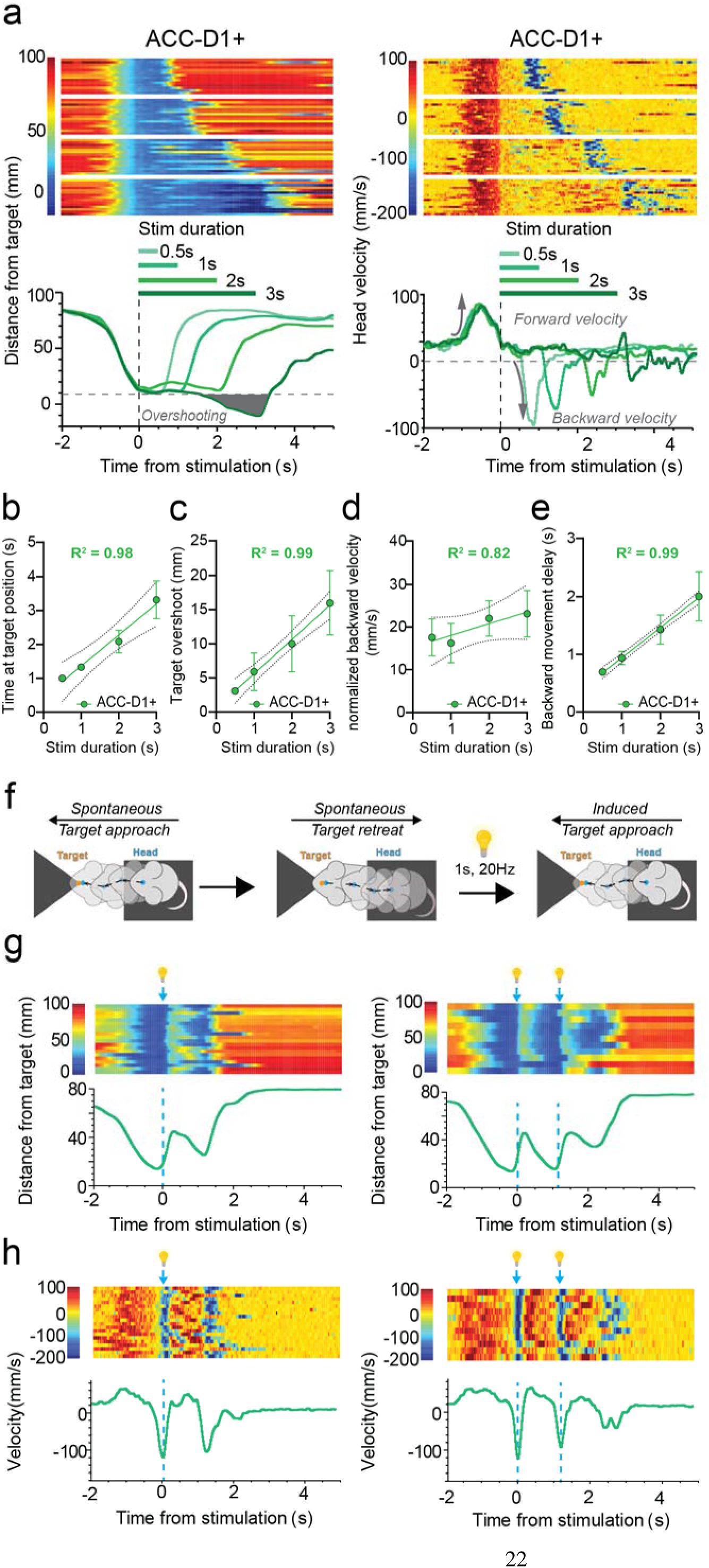
Activation of ACC-D1+ projections can cause overshooting and forward movement depending on timing of stimulation. (**a**) The effect on the approach behavior depends on stimulation duration. (Left) Longer stimulation (3 s) can result in target overshooting. (Right) Backward movement is delayed until the end of stimulation. (**b**) Different stimulation durations predict time at target position at 20 Hz. (n = 11 mice, R^2^ = 0.98). (**c**) Stimulation of ACC-D1+ also causes target overshoot (n = 11 mice, R^2^ = 0.99). (**d**) Backward velocity as a function of different stimulation durations (n = 11 mice, R^2^ = 0.82). (**e**) Backward movement delay for different stimulus lengths at 20 Hz (n = 11 mice, R^2^ = 0.99). (**f**) Illustration of stimulation upon returning to the starting position. A 1s light stimulation at 20Hz is delivered upon a spontaneous target retreat, causing an induced target approach. (**g**) Multiple stimulations can be triggered and induce repeated forward approach. (Left) Distance to target upon one stimulation upon retreat. (Right) Distance to target upon two stimulations upon retreat. (**h**) (Left) Velocity measurement when the mouse is stimulated upon retreat once. (Right) Velocity measurement when the mouse is stimulated twice.

The stimulation effect could not be explained by increased reward value at the target. At effective frequencies (10, 20, 40 Hz), the stimulation can actually induce the mouse to drop the food or skip its collection, and simply persist in moving forward for the duration of the stimulation. When this happens, mice would usually return to the starting position and approach the reward again once the stimulation has terminated. If they did not drop the food, the mice would hold their food and start eating once they returned to their starting position. Such a phenomenon is also observed in trials without stimulation, indicating that the effect of the stimulation has no relation to reward consumption.

We then tested whether the actual movement kinematics are determined by the current behavioral state by changing the timing of stimulation to a different phase of the behavior. Instead of stimulating when the mouse reaches the target, we delivered stimulation during target retreat. At this point, optogenetic stimulation (20 Hz, 1s) of ACC-D1+ projections generates a second target approach followed by a spontaneous retreat (Fig. 2f). Strikingly, we can use this stimulation protocol to repeat the forward target approach whenever the mouse has retreated to the start position (Fig. 2g, h).

These observations indicate that the activation of ACC-D1+ neurons does not generate a stereotyped action. Rather, it appears to generate a top-down command that specifies a spatial goal target, and the actual behavioral trajectory can vary depending on the current behavioral state. When the animal is already at the target, it will result in fixation at the target location for the duration of the stimulation, but when the animal is back at the starting position, it will produce forward movement to approach the target. When the stimulation duration is high, it can generate excessive overshooting in the approach behavior. Together, these findings suggest that activity in ACC-D1+ neurons represents a discrete goal location. Stimulating these neurons can produce fixation at this spatial location after target reach, overriding natural retreat behaviors. On the other hand, if triggered at the starting position, the activation of ACC-D1+ neurons causes repeated approach behaviors.

### ACC-D1+ calcium and DA signaling show opposite correlation with approach behavior

Previous studies showed that, in rats, ACC lesions impair reward approach behavior in response to predictive cues ^7,21^, suggesting that a major function of the ACC is necessary to control approach behavior. Based on our optogenetic stimulation results, we hypothesized that the activity of ACC-D1+ neurons could represent key aspects of approach behavior, such as distance to the target. To test this hypothesis, we injected a retrograde AAV expressing a Cre-dependent genetically encoded calcium indicator (jGCaMP7) into the DMS of Drd1-Cre mice and implanted a GRIN lens into the ACC of either the left or right hemisphere. We then used an endoscope to image ACC-D1+ activity during behavior in freely moving mice (Fig. 4a) ^22^. We recorded ACC-D1+ activity during the approach behavior task (Fig. 4b). The calcium imaging data were then processed using Minian to extract the location of cells and their calcium traces ^23^. Many ACC-D1+ neurons (Supplementary Fig.4a; 32.3%, 54/167) increased activity during approach (Fig. 4c). These neurons showed a strong negative correlation with distance to target (Supplementary Fig.4b, c): as distance from target decreased, calcium activity increased (Fig. 4d). A smaller group of neurons (11.4%, 19/167) were inhibited during the target approach (Fig. 4e) and their activity was positively correlated with distance to target (Fig. 4f).

**Fig. 4.**
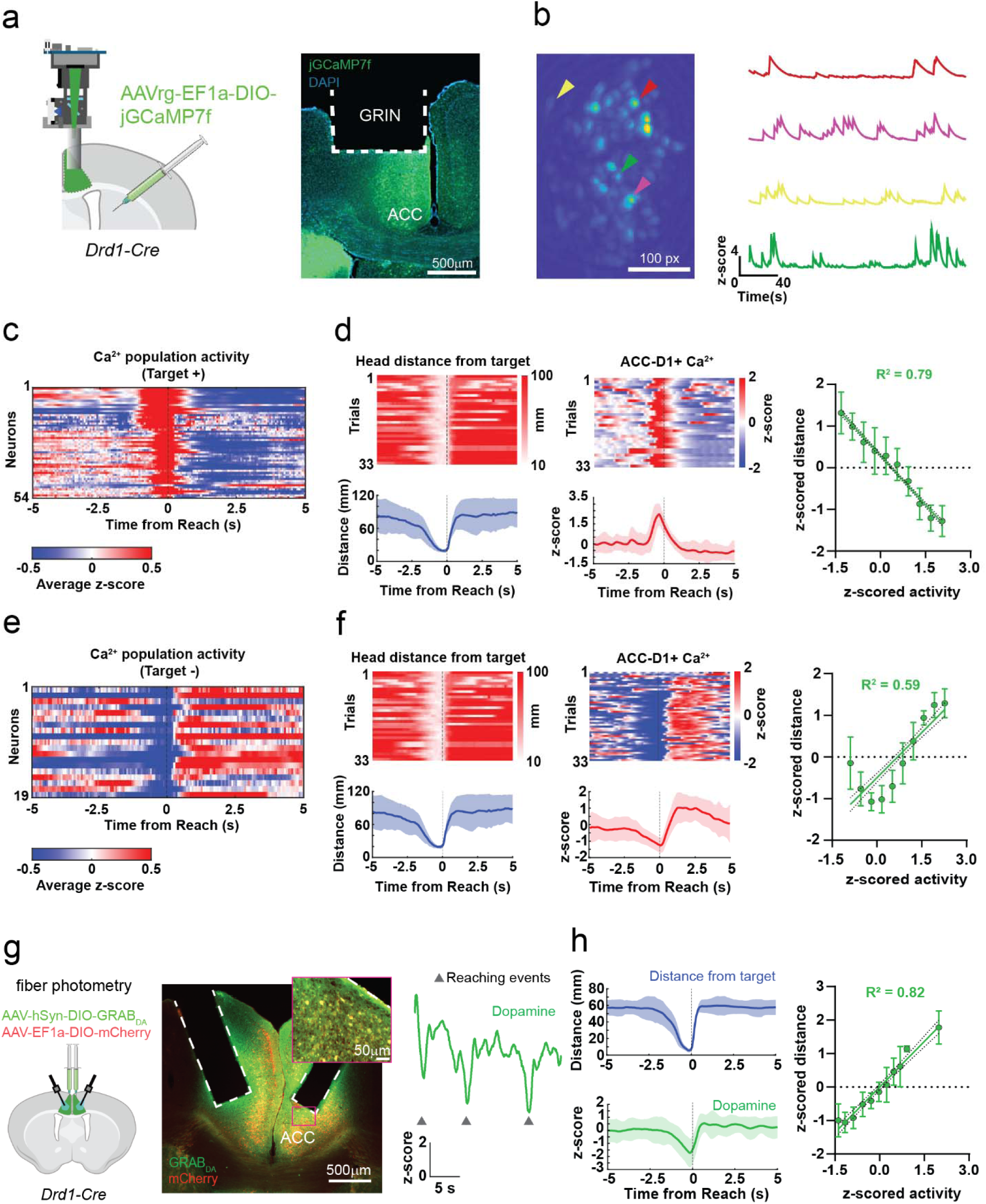
ACC-D1+ activity and ACC DA signaling during approach behavior. (**a**) (Left) Retrograde targeting of ACC-D1+ neurons using AAVrg-EF1a-DIO-jGCaMP7f injection in the DMS and positioning a 1x4 mm GRIN lens in either the left or right ACC. (Right) Representative image of a mouse brain in which, with the GRIN lens above the ACC-D1+ neurons expressing jGCaMP7f. (**b**) (Left)Maximum projection of an example field of view during calcium imaging. (Right) Example traces of Ca^2+^ activity from 4 neurons. (**c**) ACC-D1+ neurons that increase firing during approach (54/167 cells, 32.3% from 4 mice). Their activity is negatively correlated with distance to target. (**d**) (Left) Trial-by-trial raster plot and average head distance from target with an example of Ca^2+^ signals from ACC-D1+ neurons that increase firing during approach behavior. (Right) Negative correlation between neural activity with head distance from target (Ca^2+^ activity increases as the distance from target decreases) (n = 54 cells from 4 mice, R^2^ = 0.79, F-test: F (1, 538) = 2052, p < 0.0001). (**e**) Ca^2+^ activity of ACC-D1+ neurons that increases during approach behavior (19/167 neurons, 11.4%). (**f**) (Left) Examples of the trial-by-trial basis of an inhibited ACC-D1+ neuron and the corresponding distance from the target. (Right) Correlation analysis shows a positive linear correlation between calcium activity and distance from the target (19 cells from 4 mice, R^2^ = 0.59, F-test: F (1, 188) = 268, p < 0.0001). (**g**) (Left) Schematic representation of surgical procedure and fiber placement for fiber photometry experiments in Drd1-Cre mice. (Center) An example of the anatomical confirmation of GRAB_DA_ and mCherry co-expression and fiber placements. (Right) Example trace showing decreased GRAB_DA_ signal when the mouse approaches the target. (**h**) (Left) Peri-event histograms of distance from the target and DA activity both referenced to the reaching events. (Right) Positive correlation between distance from target and DA activity across all mice (8 hemispheres from 4 mice, R^2^ = 0.82; F-test: F (1, 86) = 397, p < 0.0001).

Since cortical DA is known to be important for directional movement^24^ and we found that ACC-D1+ neurons are critical for target approach and forward movement, we then measured DA dynamics in the ACC during approach behavior. We expressed a Cre-dependent GRAB-DA sensor together with a Cre-dependent fluorescent reporter (mCherry) in the ACC of Drd1-Cre mice to measure DA signaling during behavior using in vivo fiber photometry (Fig. 4g, left and center). We found that approach behavior was accompanied by a decrease in DA signaling (Fig. 4g, right). This was observed in all the recorded mice independently of the hemisphere recorded (Supplementary Fig.5). On average, we found ACC DA is positively correlated with distance to target, decreasing as the mouse approaches the target (Fig. 4h). These data show a negative correlation between DA signaling and ACC-D1+ activity, suggesting that DA may normally suppress ACC-D1+ activity. During approach behavior, DA decreases as ACC-D1+ neurons increase their activity.

**Fig. 5.**
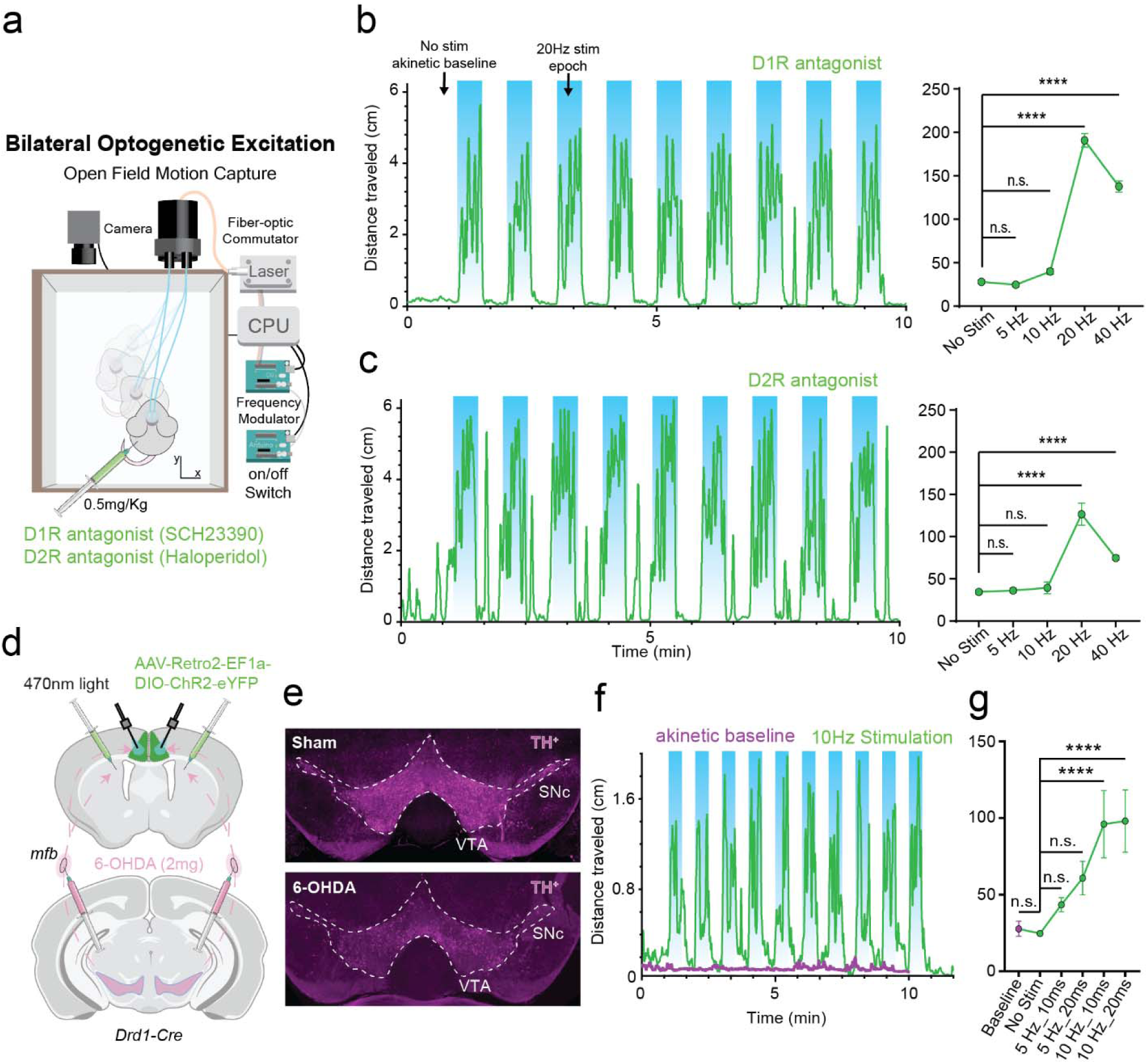
Rescue of akinesia by optogenetic activation of ACC-D1+ neurons. (**a**) Illustration of open-field test in which akinesia was induced using dopamine antagonists targeting D1 or D2 receptors. Optogenetic stimulation of ACC-D1+ neurons was then performed to restore movement using a one-minute on/off design. (**b**) (Left) Example of a mouse injected with D1R antagonist SCH23390 in which the akinesia during the “No Stim” epochs is rescued by light stimulation. (Right) Parametric quantification of the rescue effect at different stimulation frequencies. The most effective frequency for rescue is 20 Hz. One-way ANOVA for SCH23390 injected mice (n = 10 mice showed a significant main effect of stimulation. F (4,75) = 295.2, ****p < 0.0001; Tukey’s multiple comparison test is reported in the figure. (**c**) (Left) Example of a mouse injected with D2R antagonist Haloperidol in which the akinesia during the “No Stim” epochs is rescued by light stimulation. (Right) Parametric quantification of the rescue effect at different stimulation frequencies. One-way ANOVA for Haloperidol injected mice (n = 10 mice), F (4,75) = 51.71, ****p < 0.0001; Tukey’s multiple comparison test reported in the figure. (**d**) Schematic representation of bilateral 6-OHDA lesions to deplete DA and optogenetic preparation for selective excitation of ACC-D1+ neurons after DA depletion. (**e**) Representative example of reduced tyrosine hydroxylase staining (TH) in the VTA and SNc after 6-OHDA injections. (**f**) Example distance trace of a Drd1-Cre mouse with 6-OHDA lesion showing an akinetic baseline in the absence of optogenetic stimulation (purple trace) and rescue of locomotor activity using a 30-second on/off light stimulation protocol at 10 Hz. (**g**) Parametric quantification of the rescue effect using different light stimulation parameters in 6-OHDA injected Drd1-Cre mice (n = 7-9 mice). One-way ANOVA, F (5, 65) = 12.09, ****p<0.0001; Tukey’s multiple comparison test reported in the figure.

### Selective activation of ACC-D1+ neurons can rescue akinesia induced by dopamine antagonism or dopamine depletion

We found that optogenetic stimulation of ACC-D1+ projections induces a forward movement toward a target location. A target can be a stable food platform or some desired location in space in front of the animal. For instance, during locomotion, we can set environmental targets that are constantly updated ^25^. We hypothesize that optogenetic excitation of ACC-D1+ neurons could also play such a role in natural locomotion. This hypothesis predicts that activation of these neurons can also generate locomotion in an open arena devoid of specific physical targets, like a platform and specific rewards. To test this hypothesis, we used an open-field test while tracking the mouse movement using a Blackfly camera and the Bonsai software (Supplementary Fig.6a) ^26^. In each session, ACC, ACC-D2+, or ACC-D1+ neurons were stimulated every other minute at a specific frequency (5, 10, 20, or 40 Hz). We found that non-specific stimulation of ACC neurons and circuit-specific modulation of ACC-D2+ projections did not produce any effect on the mouse locomotion, whereas stimulation of ACC-D1+ neurons significantly increased locomotion. This locomotor effect is also frequency-dependent, with 20 Hz showing the most significant effect in the ACC-D1+ group (Supplementary Fig.6b,c).

In Parkinson’s disease, dopamine depletion can dramatically impair self-initiated actions and locomotion. Our results suggest that the ACC-D1+ neurons might be a novel treatment target for rescuing Parkinsonian akinesia. We first tested if stimulation could reverse akinesia induced by dopamine antagonism using intraperitoneal injections (i.p.) of D1 and D2 antagonists. Injection of the D1 antagonist SCH23390 (i.p. 0.5 mg/kg) or the D2 antagonist haloperidol (i.p. 0.5 mg/kg) caused akinesia. Following injections, we found that activation of ACC-D1+ neurons rescued akinesia induced by D1 antagonism (Fig. 5a,b) or by D2 antagonism (Fig. 5c).

We then tested the efficacy of ACC-D1+ stimulation in reversing akinesia induced by permanent DA depletion in 6-OHDA-lesioned mice, a common animal model of Parkinsonian akinesia. We expressed a retrograde AAV expressing a Cre-dependent ChR2 into the DMS of Drd1-Cre mice to excite ACC-D1+ neurons (Fig. 5d). 2mg of 6-OHDA was able to cause DA depletion, confirmed by tyrosine hydroxylase (TH) staining (Fig. 5e). Bilateral DA-depleted mice are characterized by an akinetic baseline with virtually no self-initiated movements. But this akinesia can be rescued by optogenetic stimulation of ACC-D1+ projections (Fig. 5f). We found that the rescue is most effective at 10 Hz stimulation with either 10 or 20 ms pulse width (Fig. 5g).

## Discussion

We identified and described the behavioral function of a previously unexplored population of ACC neurons expressing D1 receptors and projecting to the DMS (Fig. 1). Stimulation of these ACC-D1+ neurons produced targeted approach behavior that varies according to a specific target location. When stimulation was delivered when the mouse head was close to the target, it prolonged the duration of stay at the target and prevented backward retreat movement (Fig. 2a-f). Time spent at the target varied linearly as a function of stimulation duration, but longer stimulation durations (e.g., 3 s) can also cause an overshoot of the target. When stimulation was delivered when the mouse moved back to the starting position, however, it reliably generated forward approach behavior to the target (Fig. 3g,h). These results suggest that the activation of ACC-D1+ neurons can generate a top-down command that specifies a particular target location.

Regardless of the starting position, the activation will generate flexible approach behavior that terminates in the specified location. Strikingly, our stimulation results appear to be the opposite of symptoms in patients with ACC damage, in particular, dysmetria and reduced movement amplitude ^15^.

Thus, our results suggest that ACC-D1+ activity may encode a specific spatial goal location directly in front of the mouse. This is in agreement with previous work in rats implicating the ACC in approach behavior ^7,27^. However, in our study, the stimulation effect appears to be circuit-specific, as it is selective to the ACC-D1+ neurons that project to the DMS. By contrast, circuit-specific stimulation of ACC-D2+ neurons or non-selective stimulation of ACC neurons projecting to the DMS did not produce significant effects (Fig. 2 and Supplementary Fig.1). This observation suggests that ACC neurons that do not express D1 receptors may have different functions. Activation of other ACC neuronal populations that do not express D1 receptors may even suppress the forward approach, which may explain the lack of effects in our experimental paradigm when we did not limit channelrhodopsin expression to ACC-D1+ neurons.

Our results using calcium imaging and DA sensors (Fig. 4) revealed that the approach behavior is associated with an increase in calcium influx in ACC-D1+ neurons and a decrease in DA level within the ACC, suggesting that reduced DA signaling has a disinhibitory effect on ACC-D1+ neurons. This finding, while counterintuitive given the known role of D1 receptors in enhancing neuronal excitability, is in agreement with in vitro experiments that show a net inhibitory effect of DA or D1R agonist application on ACC neurons ^28^. This inhibitory effect was due to reduced glutamatergic signaling through AMPA receptors, but the underlying mechanisms for such effects remain unclear. Additional studies will be needed to understand the complex modulatory role of DA signaling in the ACC and its effects on behavior.

As ACC-D1+ neurons strongly project to the DMS, this corticostriatal circuit is expected to play a key role in approach behavior and locomotion. It has also been shown that striatal fast-spiking interneurons (FSIs), which also receive direct cortical projections, are critical for continuous pursuit behavior ^29^. ACC-D1+ neurons projections to FSIs may be responsible for generating the commands for target pursuit, as the FSI circuit within the striatum can convert distance to target representations to velocity commands in movement. Interestingly, in PD, akinesia can sometimes be relieved by visible targets in front of them ^30,31^. Since visual cortex input to the ACC can induce a motor response ^32^, it is possible that normally the ACC-D1+-DMS circuit is critical for targeted movements guided by a visible target.

Previous research has also shown that activation of D1+ neurons (direct pathway) in the DMS can rescue Parkinsonian symptoms like akinesia ^11^. Parkinson’s disease is associated with DA depletion in the cortex and basal ganglia, but traditional work has mostly focused on the basal ganglia. Given the robust effects of ACC-D1+ stimulation in approach behavior, we also attempted to rescue Parkinsonian akinesia using optogenetic stimulation. We induced complete akinesia using either DA antagonists or bilateral 6-OHDA lesions of DA neurons (Fig. 5). We were able to generate normal locomotion immediately upon stimulation of ACC-D1+ neurons. The immediate rescue of akinesia at relatively low stimulation frequencies (e.g. 20 Hz) suggests a treatment strategy that is distinct from conventional high frequency (>100 Hz) deep brain stimulation in the basal ganglia structures. While our findings demonstrate acute effects of ACC-D1+ stimulation, however, the long-term impact of chronic stimulation remains unexplored. In conclusion, our findings highlight the critical role of ACC-D1+ neurons projecting to the DMS in orchestrating targeted approach behavior and rescuing Parkinsonian akinesia in mice. By demonstrating that selective activation of these neurons induces flexible, location-specific movements and rescues volitional movement in akinetic mice, we uncover a novel corticostriatal circuit for volitional action. These results not only advance our understanding of the ACC’s contribution to motor control but also suggest a promising circuit-based therapeutic target for Parkinson’s disease.

## Materials and Methods

### Animals, housing, and genotyping

All mice were used in accordance with the Institutional Animal Care and Use Committee (IACUC) and the Duke Division of Laboratory Animal Resources (DLAR) oversight (IACUC Protocol Numbers A147-17-06 and A263-16-12). All mice were housed 16 (H) x 16 (W) x 25 (L) cm cages under typical day/night conditions of 12-hour cycles. All male and female Wild type C57BL/6J (Stock #000664), transgenic Drd1-Cre (B6;129-Tg(Drd1-cre)120Mxu/Mmjax; Stock #37156) or Drd2-Cre (Tg(Drd2-cre)ER43Gsat/Mmucd; Stock #017268-UCD) mice (3-5 months old) were handled for 5-10 min a day for a week to allow them to get used to the operator. After this time, animals were food-restricted for 3-5 days until they reached 85-90% of their initial body weight. The target weight was maintained stable by daily feeding them, with 1.5/2 g of home chow (Purina Rodent Chow 5053), after training. The number of mice used in each experiment is indicated in the figure legends.

### Viral injection procedure

Mice were anesthetized with 2.0 to 3.0% isoflurane mixed with 0.60 L/min of oxygen for surgical procedures and placed into a stereotactic frame (David Kopf Instruments, Tujunga, CA). Meloxicam (2 mg/kg) and topical bupivacaine (0.20 mL) were administered before incision. All the injections were performed using a microinjector (Nanoject 3000, Drummond Scientific) at a rate of 1 nL/s. For details about the injected viruses and coordinates for implants and injections, please refer to the specific description for each experimental procedure.

### Optogenetic experiments

For optogenetic experiments 200nl of AAV5-EF1a-DIO-hChR2-eYFP (Addgene - 35509) were injected in the ACC (AP: +0.9, relative to bregma; ML: ±0.12, relative to midline; DV: 1.1, relative to brain surface) and 200nl of an AAV(Retro2)-EF1a-Cre-WPRE (Duke Core - pKD134-Retro2) virus were injected in the DMS (AP: +0.5, relative to bregma; ML: ±1.4, relative to midline; DV: 2, relative to brain surface) of WT mice. Drd1-Cre and Drd2-Cre mice were injected with an AAV(Retro2)-EF1a-DIO-hChR2-eYFP (Duke Core - pKD3-retro2) in the DMS (AP: +0.5, relative to bregma; ML: ±1.4, relative to midline; DV: 2, relative to brain surface). Custom-made optic fibers (2 - 3 mm length below ferrule, >80% transmittance, 105 μm core diameter) were then implanted directly above the ACC at an angle (AP: +0.5 with respect to bregma, ML: 1.1 with respect to bregma, DV: 1.3 from the brain surface; 25°). Fibers were secured in place with dental acrylic adhered to skull screws. Mice were allowed to recover for three weeks after surgery before experimentation.

### Calcium imaging and fiber photometry

For calcium imaging experiments, a total of 500nl of AAVrg-Syn-FLEX-jGCaMP7f (Addgene - 104492) were injected monolaterally in the DMS (AP: +0.2/+0.8, relative to bregma; ML: +1.4, relative to midline; DV: 2.0, relative to brain surface). Then, a 1x4 mm gradient-index (GRIN) lens (Inscopix - 100-004113) was slowly implanted above the ACC (AP: +0.5, relative to bregma; ML: +0.5, relative to midline; DV: 1.0, relative to brain surface) to allow the live imaging of ACC-D1+ neuronal activity in freely moving animals performing a reaching task. The lens was secured to implanted cranial screws with dental cement and covered with Kwik-Sil silicone elastomer (World Precision Instruments) to protect the surface of the lens. Approximately 4 weeks later, fluorescence was checked using a UCLA miniscope designed to hold a relay lens (1.8 x 4.3 mm, Edmund Optics)(*22*), and mice were base-plated to select the preferred field of view. The analysis was performed using Minian v1.2.1(*23*).

For fiber photometry experiments, total 150nl of 4:1 mixture of AAV9-hSyn-DIO-GRAB_DA3m_ (Biohippo – PT-4719) and AAV5-EF1a-DIO-mCherry (UNC Vector Core) were injected bilaterally in the ACC (AP: +0.9, relative to bregma; ML: ±0.12, relative to midline; DV: 1.2, relative to brain surface). Optic fibers (4 mm length below ferrule, 400 μm core diameter, 0.5 NA; RWD – R-FOC-BL400C-50NA) were implanted in the ACC at an angle (AP: +0.9 with respect to bregma, ML: 1.1 with respect to bregma, DV: 1.3 from the brain surface; 25°) to allow live imaging of dopamine within the ACC. The fibers were secured to the skull using dental cement, and approximately 2 weeks later, fiber photometry data was collected at 60 Hz using a fiber photometry system (RWD R280) in freely moving mice during a reaching task. 470 nm and 560 nm excitation wavelengths (50 μW) were used, respectively, for imaging GRAB_DA3m_ and mCherry. For data preprocessing, we used the software accompanying the photometry system. Briefly, baseline correction was performed for both channels using partial least squares regression (coefficient β=8). Then, the mCherry signal was used as a control for motion correction due to its fluorescence being independent of dopamine. We fitted the mCherry signal to the GRAB_DA3m_ signal using robust least square (fitted control). ΔF/F is computed as (F-F1)/F0, where F, F1 and F0 denote GRAB_DA3m_ signal, fitted control, and median of raw GRAB_DA3m_ signal, respectively. Then, the ΔF/F was used for calculating the z-score together with the mean and standard deviation of each recording session.

### Reaching task

Two platforms were designed to stand one in front of the other at a regular distance. One platform held Bio-Serv 14 mg Dustless Precision Pellets (Bio-Serv, NJ, USA) and the other host the mouse. Mice were habituated to the platforms until they regularly reached the food platform to retrieve a pellet. For all the experimental conditions, a Blackfly camera (Flyr System, BFS-U3-04S2M-CS) was installed on top (or on the side) of the platforms to record the mouse behavior. For optogenetic stimulations, a webcam (Logitech C270) was positioned on the side of the platforms to trigger a laser stimulation based on the mouse approaching the platform or withdrawing from it. In this case, a 473 nm DPSS laser output (Shanghai Laser & Optics) was set at a power output of ∼7 mW using an analog power console (PM120VA, ThorLabs). Behavioral data, video timestamps, and TTLs were synchronized, sent and recorded using Blackrock Cerebrus recording system (Blackrock Microsystems) interfaced with Matlab. Experiments were performed by testing either a 1 second stimulation at different frequencies (5-10-20-40 Hz), or one frequency (20Hz) for different durations (0.5-1-2-3 seconds) triggered upon the mouse reaching the platform. Moreover, we set a multiple stimulation protocol upon mouse withdrawal from the platform using a 1 second laser stimulation at 20Hz. For imaging experiments, the behavioral camera and the imaging system were synchronized using Blackrock Cerebrus recording system (Blackrock Microsystems).

### Open Field experiments

Open field tests were conducted using either a stimulation every other minute or a stimulation every other 30 seconds. In both cases, WT and D1-Cre mice expressing ChR2 in ACC_→DMS_ neurons, were stimulated using 473 nm DPSS laser (Shanghai Laser & Optics) set at a power output of ∼7 mW. The Bonsai software was used to automatically detect and record at 50 frames/second through a Blackfly camera (Flyr System, BFS-U3-04S2M-CS) (*25*). The x and y coordinates for the center of mass of the mouse shape and trigger the optogenetic stimulation at the desired frequency through an Arduino Duo. To achieve short latency control of the laser, we used two Arduinos. The first Arduino served as a data acquisition device to receive digital output signals from the computer. The second Arduino modulated frequency output from the laser using transitor-transitor logic (TTL) pulses. The session began, after the mice were acclimated for 30 min to the new room. In the rescue experiments of akinesia, mice received IP injection at 0.5 mg/Kg of either a D1 antagonist (SCH23390, abcam #ab120597) or a D2 antagonist (Haloperidol, Tocris #0931/100) and experiments started 5-10 minutes after IP injection.

### 6-OHDA lesion

To generate bilateral PD mice, we prepared a solution of 2.6 mg/mL of 6-OHDA (Tocris 2547) with 1 mg/mL of L-Ascorbic Acid (Tocris 4055) in Normal Saline 0.9%. Drd1-Cre mice were injected with AAV-Retro2-EF1a-DIO-hChR2-eYFP (Duke Core - pKD3-retro2) in the DMS (AP: +0.5, relative to bregma; ML: ±1.4, relative to midline; DV: 2, relative to brain surface); 3 weeks later, 800 uL of 6-OHDA was bilaterally injected in the medial forebrain bundle (mfb, AP: -0.95, relative to bregma; ML: ±1.15, relative to midline; DV: 4.5 relative to brain surface) using a microinjector (Nanoject 3000, Drummond Scientific) at a rate of 2 nL/s. At the end of the injection, the needle was kept in place for 5 minutes before extraction to ensure sufficient uptake of the drug. Custom-made optic fibers (2 - 3 mm length below ferrule, >80% transmittance, 105 μm core diameter) were then implanted directly above the ACC at an angle (AP: +0.5 with respect to bregma, ML: 1.1 with respect to bregma, DV: 1.3 from the brain surface; 25°). Fibers were secured in place with dental acrylic adhered to skull screws. After 5-6 days, mice were placed at the center of a rectangular open field arena, as described in the open field experiment paragraph, and, after recording 10 minutes of akinetic baseline, stimulation was triggered every 30 seconds at 5 and 10 Hz with either 10 or 20 ms pulse width.

### Immunohistochemistry

After the behavioral tests were concluded, mice were anesthetized with 200 mg/kg tribromoethanol (avertin) and euthanized by perfusing with a solution made of TBS with heparin (0.1128 g Heparin ammonium salt from porcine intestinal mucosa [Sigma; H6279]) and then with 4% Paraformaldehyde (PFA). Mouse brains were then kept in 4% PFA o.n. at 4 °C. The day after brains were rinsed three times with TBS, immersed in 30% Sucrose in TBS, and stored at 4 °C until they were not floating anymore in the solution. At this time brains were included in a mixture of 30% Sucrose in TBS and Tissue Tek O.C.T. compound (frozen tissue matrix) at a 2:1 ratio and stored at −80 °C. Brains were cut into 20 or 50-μm coronal sections and stored in a 1:1 mixture of TBS/glycerol at −20 °C. Sections were washed in 1× TBS containing 0.2% Triton-X100 (TBST) and blocked in 5% NGS diluted in TBST. For the immunolabeling of ChR2, sections were incubated o.n. with a primary antibody chicken anti-GFP (1:1000; Millipore, AB16901; Aves Labs GFP 1010). For TH detection, sections were incubated o.n. with a primary antibody rabbit anti-TH (1:1000; Merck 657012). Secondary Alexa-fluorophore (488, 594) conjugated antibodies (Invitrogen) were added (1:200 in TBST with 5% NGS) for 2 h at room temperature. Slides were mounted in Vectashield with DAPI (Vector Laboratories, CA) and images were acquired on an Olympus Fluoview confocal microscope using ×20 objective at ×1.3 Zoom or using the Keyence benchtop microscope.

### Quantification and statistical analysis

The values and error bars reported in the text are the mean ± SEM, respectively. Behavioral data were analyzed with Neuroexplorer (Plexon) and MATLAB 2016b (MathWorks). Statistical tests were performed in Prism 8 (GraphPad). Two-tailed parametric tests were used. An a priori alpha level of 0.05 was used to determine significance. All the details about the statistical test used and the relevant information are reported in the figure legends.

## Acknowledgments

Illustrations were created with BioRender.com. We thank Donna Porter and Fengxia Allen for helping with mouse colony maintenance, Guozhong Yu for his help with optic fiber fabrication and brain harvesting, L. Menéndez de la Prida and M. Angeles Arevalo for the constructive discussion and feedback on the manuscript. C.E. is an HHMI Investigator. This work was supported by the Regeneration Next Postdoctoral fellowship, 4511578 to FPUS, the RYC2021-033202-I to FPUS, the PID2023-146385NA-I00 by FPUS, NINDS grant R01-NS094754 to HHY, the N.I.H. R01-NS102237 to CE, and by the joint efforts of MJFF and the Aligning Science Across Parkinson’s (ASAP) initiative grant ASAP-020607 to CE.

## Funding

Regeneration Next Postdoctoral fellowship, 4511578 (FPUS)

This work was supported by the grant RYC2021-033202-I, funded by MCIN/AEI/10.13039/501100011033 and European Union «NextGenerationEU»/PRTR» (FPUS)

This work was supported by the grant PID2023-146385NA-I00, funded by MCIU/AEI/10.13039/501100011033 and FSE+ (FPUS)

National Institute of Neurological Disorders and Stroke, R01-NS094754 (HHY) National Institute of Health, R01-NS102237 (CE)

Aligning Science Across Parkinson’s, ASAP-020607 (CE)

## Author contributions

Conceptualization: FPUS, CE, HY Methodology: FPUS, BL, JK, AF, MR, SAJ, KB

Investigation: FPUS, BL, JK, AF, MR, SAJ, KB Visualization: FPUS, CE, HY

Funding acquisition: FPUS, CE, HY Writing – original draft: FPUS, JK, CE, HY Writing – review & editing: FPUS, CE, HY

## Data and materials availability

All data are available in the main text or the supplementary materials.

