## Supplementary Figures for "Elucidating an anterior cingulate circuit for self-initiated actions and rescue of Parkinsonian akinesia"

**Affiliations**


**This PDF file includes:**

Supplementary Figure 1-6
Legend for Supplementary Movie 1, 2

**Other Supplementary Materials for this manuscript include the following:**

Supplementary Movie 1, 2

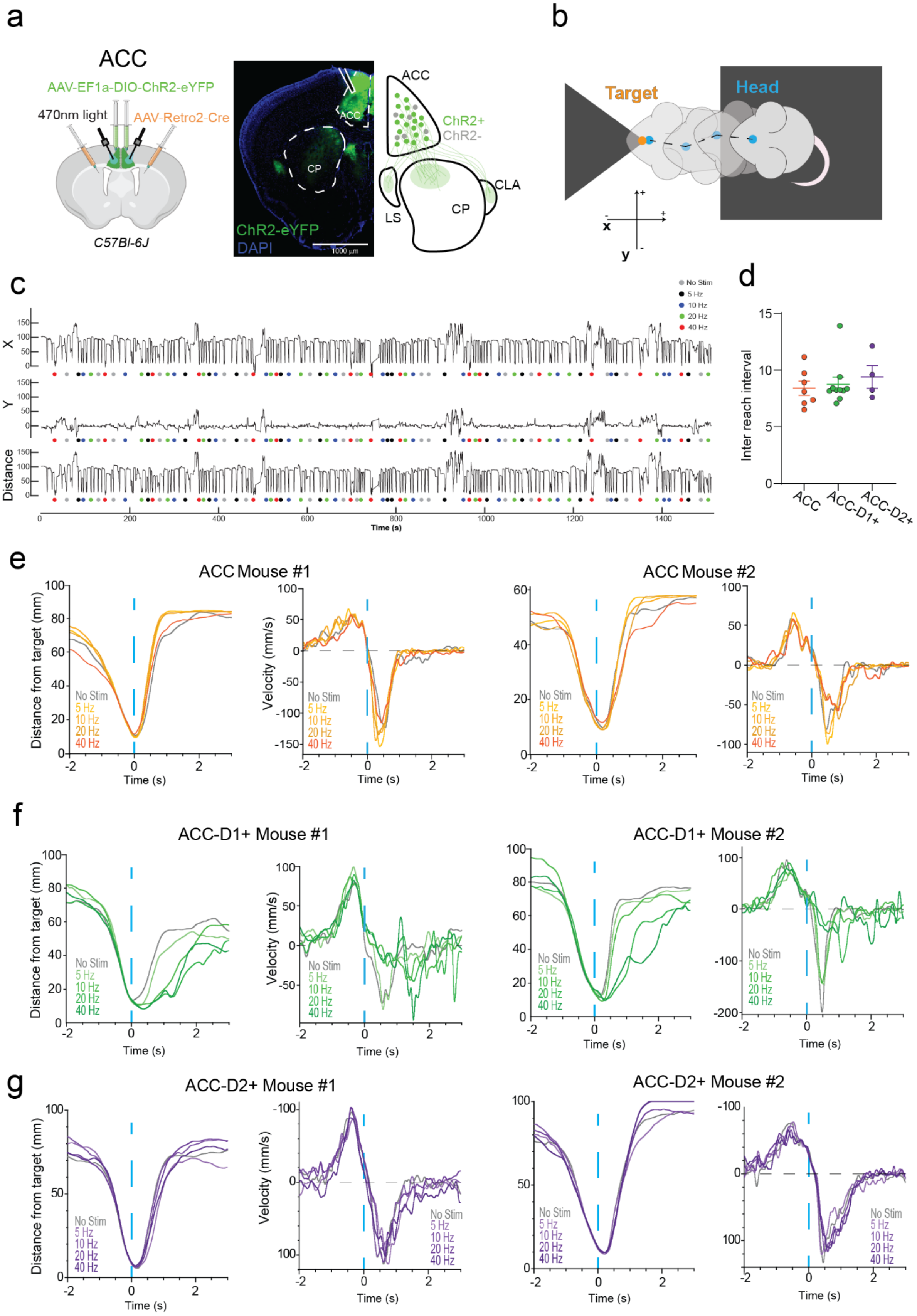
a
ACC
AAV-EF1a-DIO-ChR2-eYFP
470nm light
AAV-Retro2-Cre
C57Bl-6J
ACC
CP
ChR2-eYFP
DAPI
1000 µm
ChR2+
ChR2-
LS
CLA
b
Target
Head
x
y
c
No Stim
5 Hz
10 Hz
20 Hz
40 Hz
X
Y
Distance
Time (s)
d
Inter reach interval
ACC
ACC-D1+
ACC-D2+
e
ACC Mouse #1
ACC Mouse #2
Distance from target (mm)
Velocity (mm/s)
Time (s)
f
ACC-D1+ Mouse #1
ACC-D1+ Mouse #2
g
ACC-D2+ Mouse #1
ACC-D2+ Mouse #2

**Supplementary Figure 1. Representative examples of behavioral measures showing response to different frequencies of photo-stimulation.** (**a**) Cre-dependent ChR2 expression in the ACC circuit in Wild-type mice (C57Bl-6J). Retrograde labeling of ACC neurons that project to the DMS. LS, lateral septum, CLA, claustrum. (**b**) Schematic of mouse approach to target and spatial coordinate reference. (**c**) Example trace for head x and y coordinates and calculated head distance from target for an entire session. Dots represent the randomly selected stimulation pattern. (**d**) Inter reach interval for all groups. One-way Anova shows no difference among them $F(2,18) = 0.3646$, $p = 0.6995$. (**e**) Distance to target and movement velocity in 2 mice with non-selective activation of ACC neurons. (**f**) Distance to target and movement velocity in 2 mice with selective activation of ACC-D1+ neurons. Stimulation prolonged the time spent at the target. (**g**) Distance to target and movement velocity in 2 mice with selective activation of ACC-D2+ neurons. Stimulation at different frequencies did not produce significant effects.

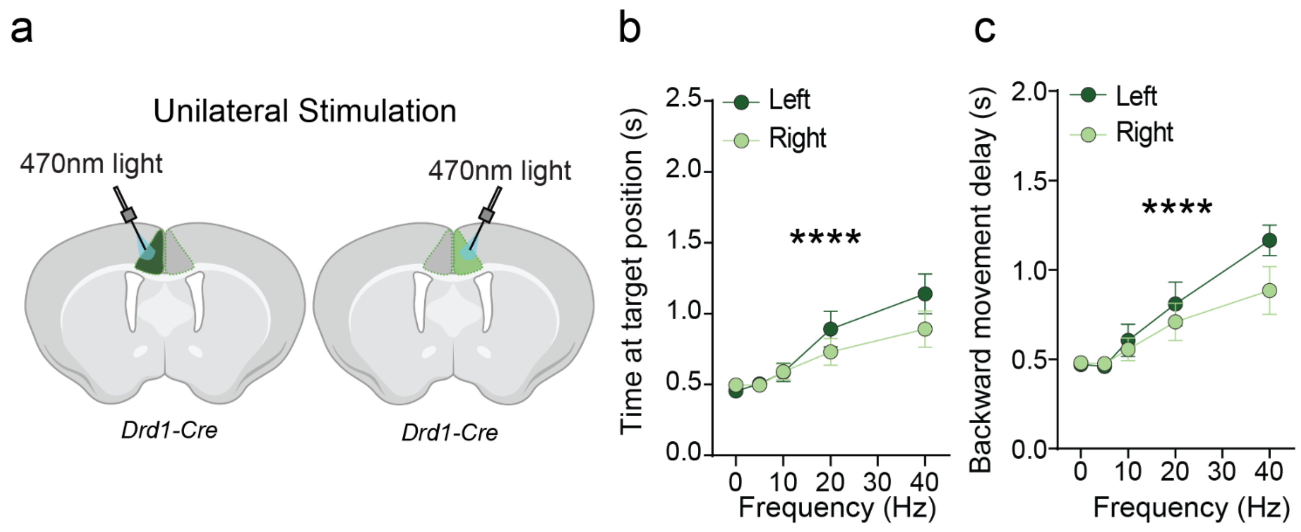


**Supplementary Figure 2. Unilateral stimulation of ACC-D1+ projections produced a weaker effect on movement.** (**a**) Illustration of unilateral stimulation of ACC-D1+ neurons. (**b**) Unilateral stimulation of ACC-D1+ projections (n = 10 mice). 2way ANOVA RM for the time spent at the target position for multiple stimulation frequencies (0, 5, 10, 20, and 40 Hz) shows a significant effect of frequency F (1.524, 27.43) = 33.25, p <0.0001; no significant effect of hemisphere F (1, 18) = 0.5879, p = 0.4532 and no interaction F (4, 72) = 2.419, p = 0.0563. (**c**) Similarly, 2-way ANOVA RM for the peak backward velocity time shows a significant effect of frequency F (2.521, 45.38) = 25.12, p < 0.0001; no significant effect of hemisphere F (1, 18) = 1.013, p = 0.3275, and no significant interaction F (4, 72) = 1.661, p = 0.1685.

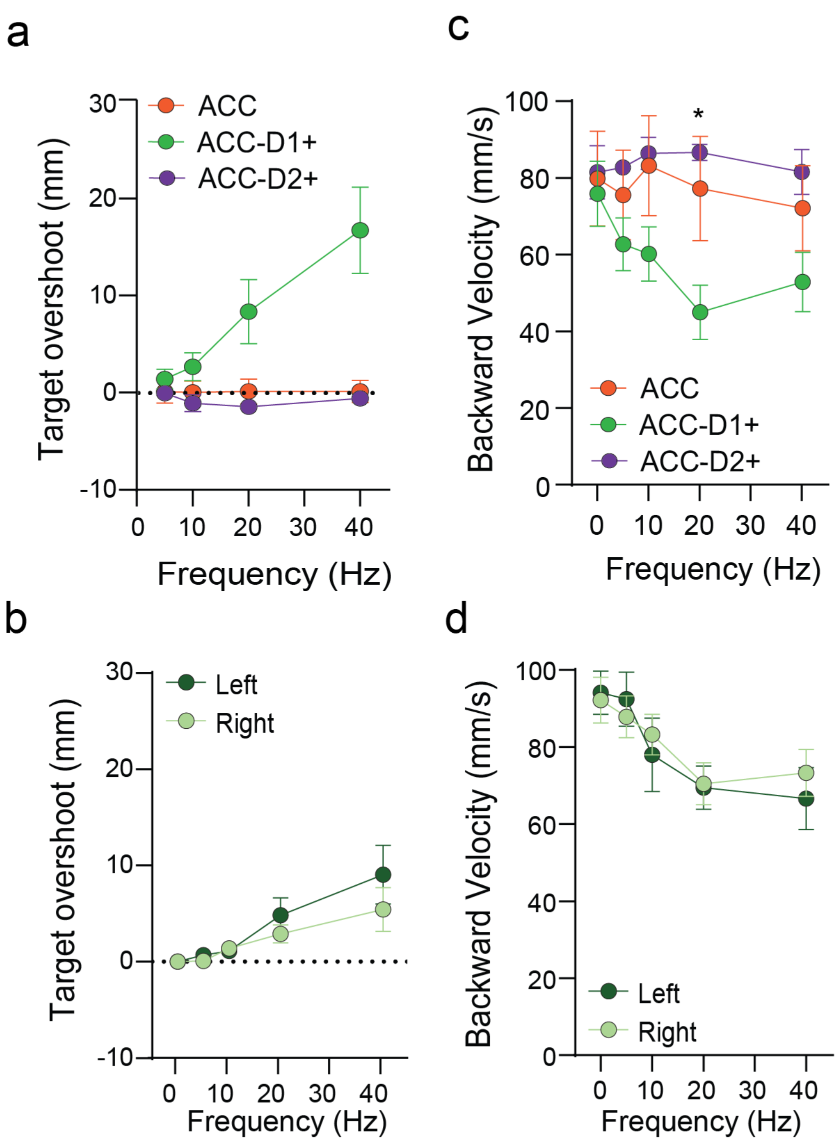


**Supplementary Figure 3. Parametric analysis of target overshoot effect.** (**a**) The head max distance from the target was normalized on the No Stim condition for all groups, ACC projection (n = 7 mice), ACC-D2+ projection (n = 4 mice), and ACC-D1+ projection (n = 10 mice). 2way ANOVA RM shows no significant effect of frequency F (1.626, 29.27) = 3.545, p = 0.0504; a significant effect of projection-specificity F (2, 18) = 4.999, p = 0.0188 and significant interaction F (6, 54) = 4.648, p = 0.0007. (**b**) Parametric analysis of peak backward velocity for all groups, ACC projection (n = 7 mice), ACC-D2+ projection (n = 4 mice), and ACC-D1+ projection (n = 10 mice). 2way ANOVA RM shows no significant effect of frequency F (2.289, 41.20) = 2.323, p = 0.1039; no significant effect of projection-specificity F (2, 18) = 0.9808, p = 0.3942 and significant interaction F (8, 72) = 2.207, p = 0.0367. (**c**) Parametric analysis of target overshoot effect. The head max distance from the target was normalized on the No Stim condition for unilateral stimulation of ACC-D1+ projection (n = 10 mice). 2way ANOVA RM shows a significant effect of frequency F (1.248, 22.46) = 13.36, p = 0.0007; no significant effect of hemispheres F (1, 18) = 0.7434, p = 0.3999 and no significant interaction F (4, 72) = 0.9379, p = 0.4471. (**d**) Parametric analysis of peak backward velocity for unilateral stimulation of ACC-D1+ projection ($n = 10$ mice). 2way ANOVA RM shows a significant effect of frequency $F(2.213, 39.83) = 8.482$, $p = 0.0006$; no significant effect of hemispheres $F(1, 18) = 0.0423$, $p = 0.8392$ and no significant interaction $F(4, 72) = 0.4004$, $p = 0.8077$.

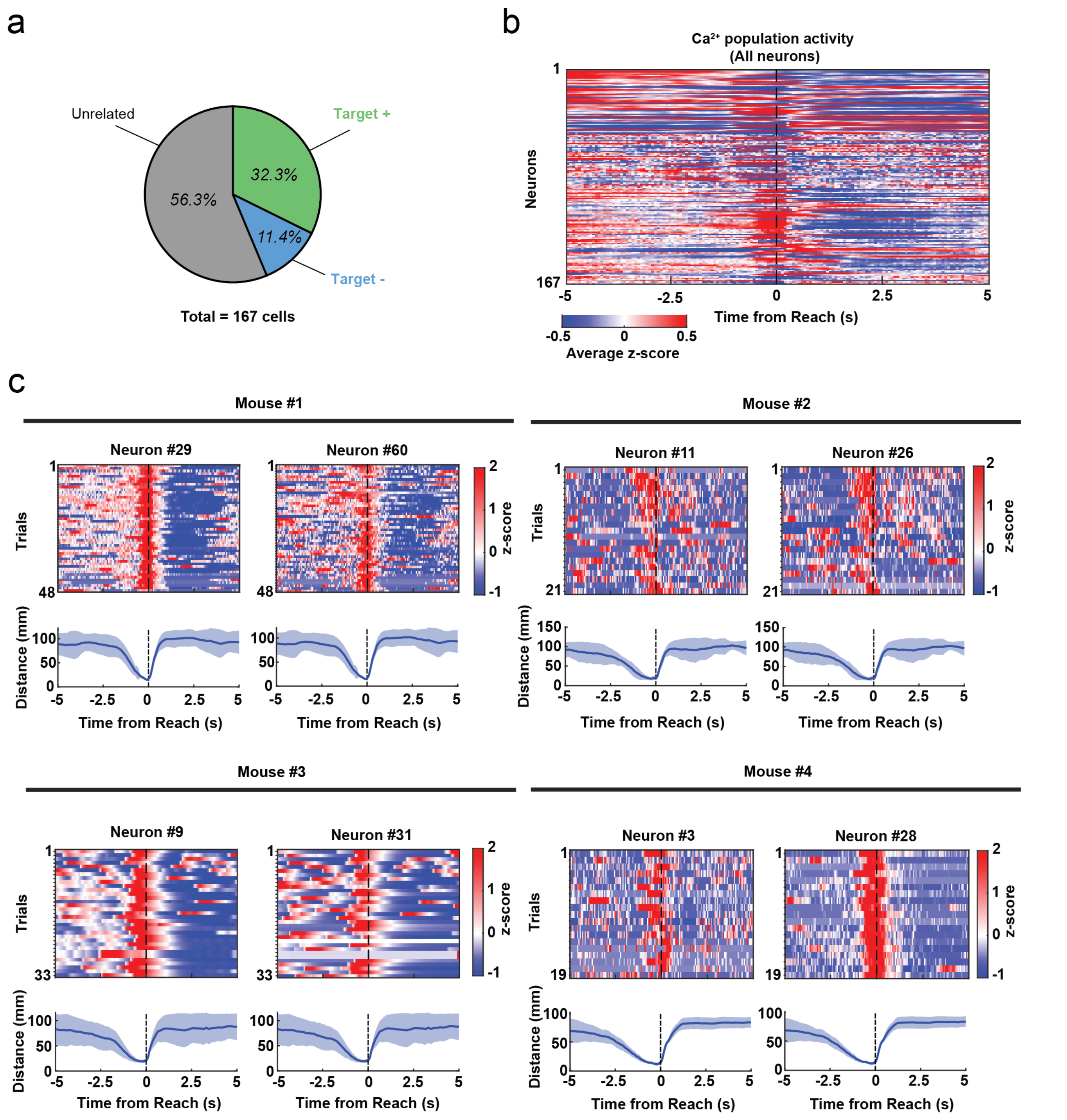


**Supplementary Figure 4. Additional examples of calcium imaging results from different types of ACC-D1+ neurons.** (**a**) The pie chart shows the relative proportions of different ACC-D1+ populations. (**b**) Calcium population activity of all neurons recorded from all the mice. (**c**) Example of two "Excited at reach" ACC-D1+ neurons for each mouse (2 mice recorded from the left hemisphere and 2 mice recorded from the right hemisphere) to show how, on a trial-by-trial basis, the inverse correlation with approach behavior remains consistent and independent of the hemispheres.

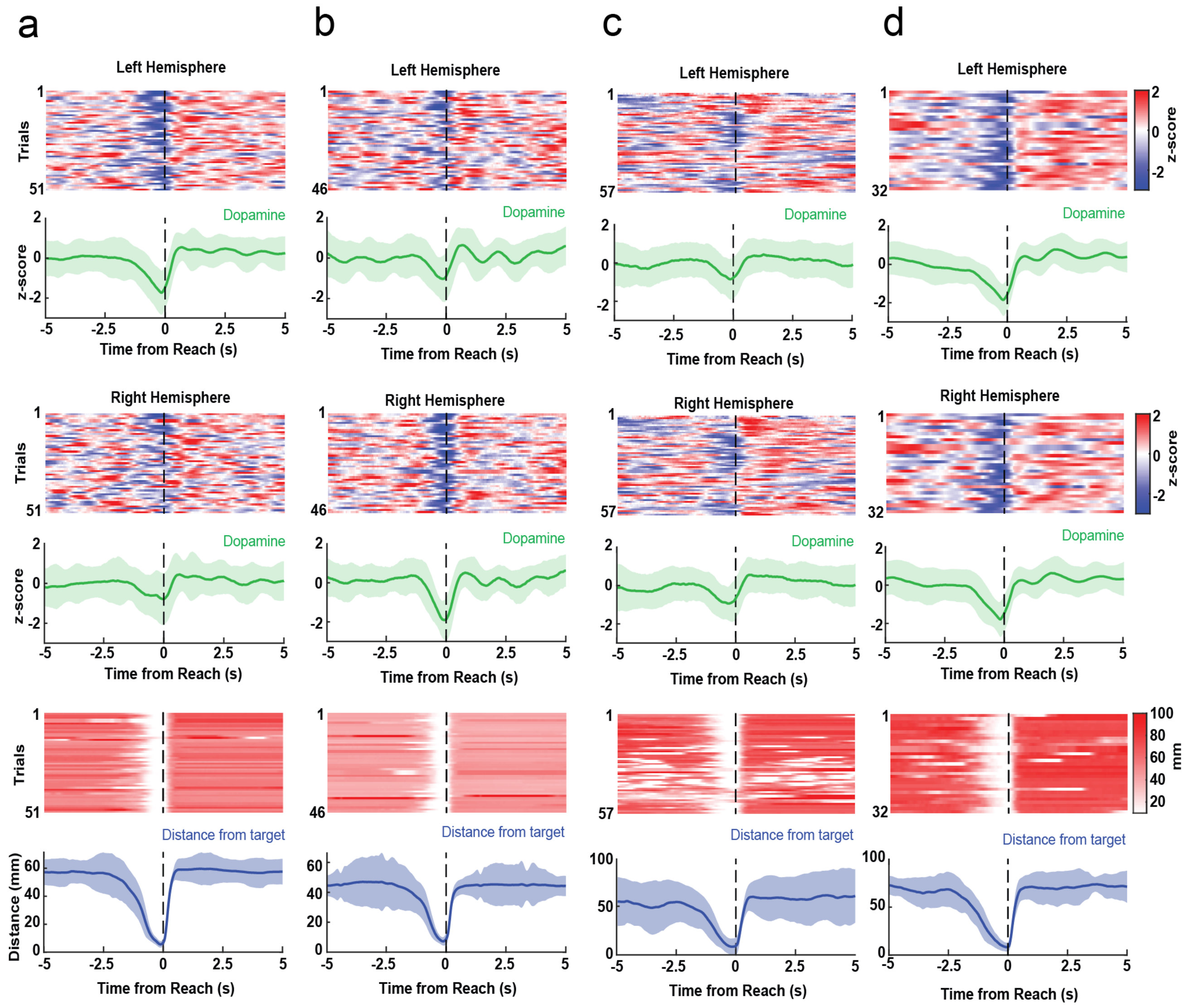


**Supplementary Figure 5. GRAB-DA sensor activity from individual hemispheres. (a-d)** Example for each of the four bilateral fiber photometry recordings of ACC-D1+ neuronal population and the corresponding distance from the target variable showing a trial-by-trial consistency of diminished dopamine level with target approach.

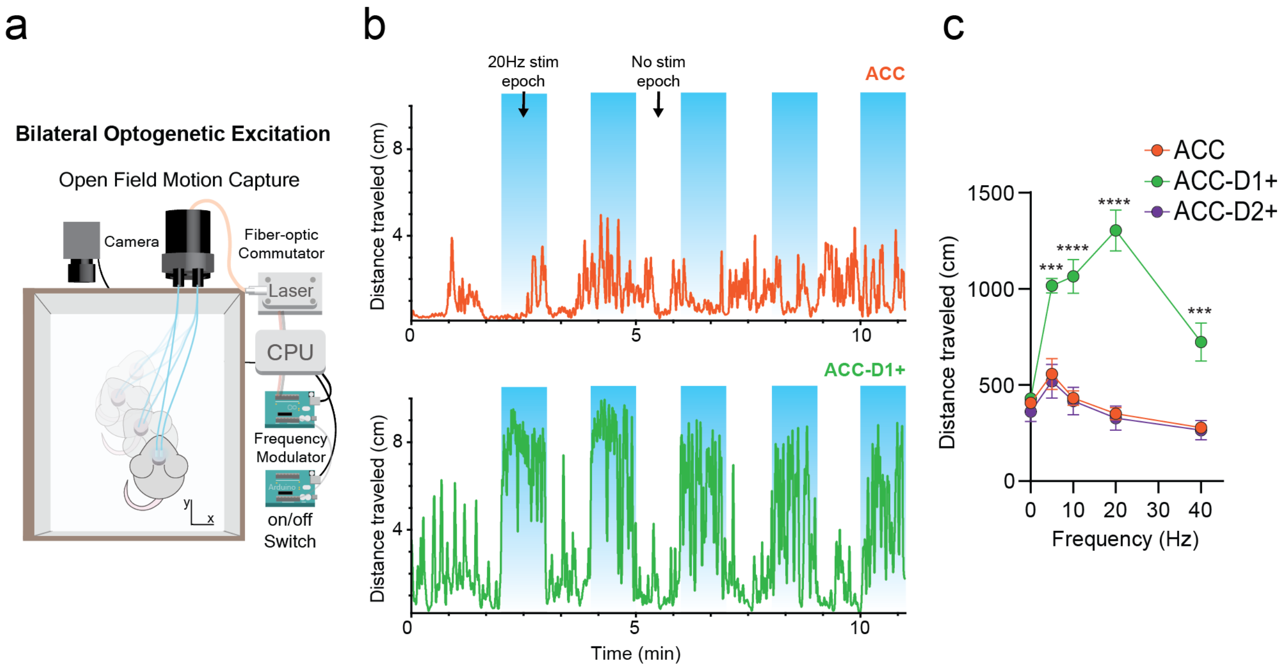


**Supplementary Figure 6. Optogenetic stimulation of different ACC neuronal populations in normal mice.** (**a**) Schematic representation of an open-field test in which either C57BL/6J (ACC), Drd2-Cre (ACC-D2+) or Drd1-Cre (ACC-D1+) mice are optogenetically stimulated with a 1-minute on/off protocol. (**b**) (Top) Example of a C57BL/6J (ACC) mouse in which the locomotor activity is not coupled to the stimulation. (Bottom) Example of a Drd1-Cre (ACC-D1+) mouse in which locomotor activity increases during the stimulation times. (**c**) Parametric quantification of distance traveled for all groups, ACC projection (n = 7 mice), ACC-D2+ projection (n = 4 mice), and ACC-D1+ projection (n = 9 mice). are optogenetically stimulated. 2way ANOVA RM: significant effect of frequency F (4, 85) = 10.47, p < 0.0001; significant effect of projection-specificity F (2, 85) = 95.47, p < 0.0001; Significant interaction F (8, 85) = 8.127, p < 0.0001. Sidak`s multiple comparison (reported in the figure) shows a significant effect between the ACC and the ACC-D1+ groups: 5 Hz (p = 0.0004), 10 Hz (p < 0.0001), 20 Hz (p < 0.0001), and 40 Hz (p = 0.0007).

**Supplementary Movie 1.**
ACC-D1+ bilateral stimulation at different frequencies

**Supplementary Movie 2.**
Bilateral ACC-D1+ stimulation in bilateral 6-OHDA mice
